# Cell cycle dynamics and window of MUTE action during the terminal division of stomata and ramification of its loss-of-function on the cell cycle of an uncommitted precursor

**DOI:** 10.1101/2022.09.25.509410

**Authors:** Daniel T. Zuch, Arvid Herrmann, Eun-Deok Kim, Keiko U. Torii

## Abstract

Plants develop in the absence of cell migration. As such, cell division and differentiation need to be coordinated for functional tissue formation. Cellular valves on the plant epidermis, stomata, are generated through a stereotypical sequence of cell division and differentiation events. In Arabidopsis, three master-regulatory transcription factors, SPEECHLESS (SPCH), MUTE, and FAMA sequentially drive initiation, proliferation, and differentiation of stomata. Among them, MUTE switches the cell cycle mode from proliferative asymmetric division to terminal symmetric division and orchestrates the execution of the single symmetric division event. However, it remains unclear to what extent MUTE regulates the expression of cell cycle genes through the symmetric division and whether MUTE accumulation itself is gated by the cell cycle. Here, we show that MUTE directly upregulates the expression of cell cycle components throughout the terminal cell cycle phases of a stomatal precursor, not only the core cell cycle engines but also checkpoint regulators. Time-lapse live imaging using the multi-color cell cycle indicator PlaCCI revealed that MUTE accumulates up to the early G2 phase, whereas its successor and direct target, FAMA, accumulates at late G2 through terminal mitosis. In the absence of MUTE, meristemoids fail to differentiate, and their G1 phase elongates as they reiterate asymmetric divisions. Combined, our work provides the framework of cell cycle and master regulatory transcription factors to coordinate a single symmetric cell division, and suggests a mechanism for the eventual cell cycle arrest of an uncommitted stem-cell-like precursor at the G1 phase.

## INTRODUCTION

Multicellular organisms grow and develop through a series of cell divisions, which coincide with the deployment of gene regulatory networks to create diverse, organized tissues. For plants, in which cells are immotile, cellular divisions and growth must be carefully coordinated in order to properly distribute specific cell types and drive tissue shape formation. As such, plants have evolved to utilize multiple environmental and developmental pathways to regulate cell cycle (Harashima and Schnittger 2010; Shimotohno et al. 2021). Importantly, accumulating evidence has shown that specific cell cycle regimes are closely integrated with the differentiation of specific cell types – allowing plants to dynamically tune cell type composition, growth, and patterning during tissue formation (Inze and De Veylder 2006; Shimotohno et al. 2021).

Core cell cycle engines are highly conserved across eukaryota and can be divided into five distinct phases (Gap 0 (G0), Gap 1 (G1), DNA Synthesis (S), Gap 2 (G2), Mitosis (M)) (Harashima et al. 2013; Johnson and Walker 1999). Each cell-cycle phase relies upon phase-specific cyclins, cyclin-dependent kinases (CDKSs), and transcription factors to drive the cell cycle forward. Temporally regulated synthesis and stability of cyclins/CDKS, as well as CDK inhibitors tune the activity of cyclin/CDK complexes to ensure the progression and timing of cell cycle phases (Harashima et al. 2013; Johnson and Walker 1999). It has been shown that plants encode a large number of core cell-cycle genes and their modifiers, likely reflecting a diverse array of cell-cycle states and context-specific modification of cell cycle dynamics (Peres et al. 2007; Riou-Khamlichi et al. 2000; Shimotohno et al. 2021; Sozzani et al. 2010).

Of particular importance to the fitness of the land plants is the development and distribution of stomata, or adjustable pores, on the leaf surface - which facilitate gas exchange and transpiration. Studies in Arabidopsis have shown that stomatal differentiation occurs through a series of stereotypical cell divisions controlled by three master basic-helix-loop-helix (bHLH) transcription factors, SPEECHLESS (SPCH), MUTE, and FAMA (Han and Torii 2016; Lau and Bergmann 2012). SPCH initiates and maintains asymmetric divisions (ACDs) of stem-like stomatal precursors called meristemoids (MacAlister et al. 2007; Pillitteri et al. 2007). After a few rounds of ACDs, MUTE drives the differentiation of a mersitemoid into a guard mother cell (GMC) and simultaneously orchestrates a single, terminal symmetric division (SCD) (Han et al. 2018; Pillitteri et al. 2007). Lastly, FAMA terminates the cell cycle in each daughter cell and completes the terminal differention of guard cells (Ohashi-Ito and Bergmann 2006).

How the stomatal-lineage bHLH proteins integrate with cell cycle machinery to switch cell cycle modes is an important question, and previsous studies have revealed several key links (Fig. 1A) (Han and Torii 2019). For instance, SPCH initiates ACDs by inducing the expression of the G1-specific D-Type Cyclin CYCD3;2 (Lau et al. 2014), which likely complex with CDKA;1 to launch the cell cycle and drive mitotic divisions (Dewitte et al. 2007; Healy et al. 2001). MUTE then disengages the SPCH-mediated ACDs by directly upregulating SIAMESE RELATED4 (SMR4), which inhibits CYCD3s but permits MUTE-induced G1 cyclin, CYCD5;1 to proceed to a terminal SCD (Han et al. 2022). An additional G1 cyclin, CYCD7;1 may also coordinate the SCD (Weimer et al. 2018). Lastly, FAMA and the Myb protein FOUR LIPS (FLP), which are directly induced by MUTE, suppress cell cycle by directly inhibiting the expression of multiple core cell cycle genes including *CDKB1;1*, thereby ensuring that the terminal symmetric division occurs only once (Hachez et al. 2011; Xie et al. 2010).

**Fig. 1.**
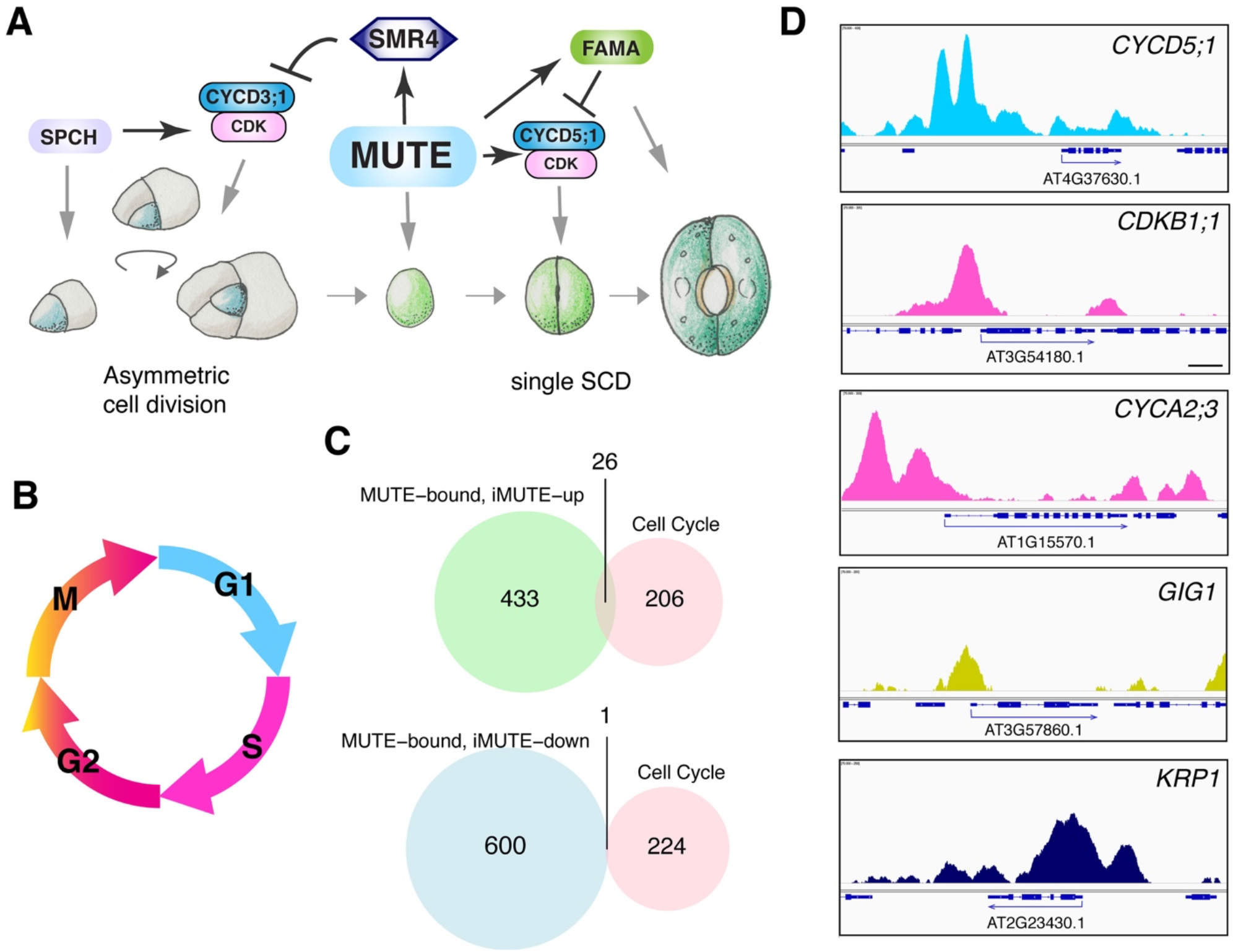
MUTE directly upregulates cell cycle regulatory genes. (A) Schematic model of how MUTE promotes GMC SCD and stomatal differentiation via cell cycle regulatory genes. (B) Phases of cell cycle color coded according to PlaCCI fluorophore expression window (phases not to scale) (C) Venn diagrams of MUTE bound & iMUTE-up (light green), MUTE bound & iMUTE-down (light blue), and a curated cell cycle gene (light pink). For gene lists, see Table S1. (D) Genome browser view of ChIP-seq profile of MUTE binding to the promoter of *CYCD5;1, CDKB1;1, CYCA2;3, GIG1*, and *KRP1*. Peaks are color coded according to their general expression window as in panel B; KRP1 is highlighted as navy blue-to show it is repressed by MUTE.

While recent efforts have uncovered several mechanisms by which the stomatal differentiation programs utilize cell cycle machinery, it remains unclear whether the cell cycle gates the expression of master regulatory stomatal-lineage bHLH proteins. This is most critical for MUTE and FAMA, in which sequential actions occur within a single round of cell cycle during the terminal SCD. Here, we show that MUTE directly upregulates a suite of cell cell cycle components throught the cell cycle phases during the terminal SCD, whereas both MUTE and FAMA exhibit temporally-restricted expression profiles that coincide with specific cell cycle phases. Furthermore, in the absence of *MUTE*, the G1 phase of dividing meristemoids becomes progressively extended as meristemoids reitrate ACDs. Together, our work highlights the cell-cycle windows during which MUTE and FAMA accumulate to coordinate a single symmetric cell division, and suggests a mechanism for the eventual cell cycle arrest of uncommitted *mute* meristemoids at the G1 phase.

## RESULTS

### MUTE directly induces a suite of cell cycle regulators throughout the cell cycle duration

Previous studies have revealed that SPCH, MUTE, and FAMA tightly regulate stomatal development through the control of both differentiation and cell cycle (Fig. 1A). Among them, MUTE must precisely coordinate the differentiation and single symmetric division of meristemoids into pairs of guard cells (Han et al. 2022; Han et al. 2018). Here, we explore the relationship between MUTE and cell cycle machinery to understand how MUTE can reliably orchestrate the shift from proliferation to a single terminal division. Notably, MUTE overexpression (iMUTE) was shown to upregulate a suite of genes involved cell cycle, cell division, and mitosis (Han et al. 2018), together representing a complete and independent cell cycle module (Fig. 1B). To address which of those genes are direct MUTE targets, we compared *iMUTE* RNA-seq (Han et al. 2018) and MUTE ChIP-seq (Han et al. 2022) data to extract a subset of cell cycle genes that are directly up-or down-regulated by MUTE (Fig. 1C). Among the 286 combined and curated cell cycle genes, 26 are both bound by MUTE and upregulated by iMUTE (Fig. 1C, Table S1). Those direct MUTE targets include SMR4, a known cell cycle inhibitor that slows down the fast proliferative asymmetric cell division of a meristemoid, as well as *CYCD5;1*, a known G1 cyclin that subsequently drives the GMC symmetric division (Han et al. 2022; Han et al. 2018).

From our analysis, we found that MUTE-direct targets include genes implicated in G1/S checkpoint control, *E2FF/DEL3* (*AT3G01330*) and *E2FC* (*AT1G47870*), which are known components of the DREAM Complex (Lang et al. 2021), as well as E2F target gene *ETG1* (*AT2G40550*) (Fig. 1, Table S1). In addition, MUTE directly upregulates the expression of the DREAM Complex components *TESMIN-LIKE CXCs (TCXs)*, including *TCX2/SOL2 (AT4G14770), TCX3/SOL1 (AT3G22760)*, and *TCX4 (AT3G04850). TCX2/SOL2* has been identified as stem cell ‘ubiquitous’ genes that likely play a role in cell divisions in diverse plant stem cell populations (Clark et al. 2019). *TCX2/SOL2* and *TCX3/SOL1* are direct SPCH-targets, and their loss-of-function mutations confer aberrant stomatal-lineage divisions and occasional misregulation of guard cell fate (Simmons et al. 2019). Our finding that they are also direct transcriptional targets of MUTE is consistent with the roles of *TCX2/SOL2* and *TCX3/SOL1* in the stomatal fate commitment process.

MUTE also directly induces genes attributed to S-phase progression, including the DNA replication gene *MCM3* (*MINICHROMOSOME MAINTENANCE 3*: AT5G46280) (Stevens et al. 2002), *TSO2 (AT3G27060)* (Wang and Liu 2006), and several A-type Cyclins (*CYCA1;1: AT1G44110, CYCA2;3: AT1G15570, CYCA3;2: AT1G47210*) that are expressed through the S-phase (Gutierrez 2009). In addition to G1, G1/S, and S-phase genes, G2-associated mitotic cyclin *CYCB2;3 (AT1G20610)* as well as CDKs *CDKB1;1 (AT3G54180)* and *CDKB2;1 (AT1G76540)* (Romeiro Motta et al. 2022) are also identified as direct MUTE targets upregulated by MUTE (Fig. 1., Table S1) Further, a gene regulating G2/M checkpoint by negatively regulating the anaphase promoting complex (APC/C), *GIG1* (*GIGAS CELL1*: *AT3G57860*), is directly bound and upregulated by MUTE (Fig. 1., Table S1). In accordance with our findings that MUTE induces machinery that largely favors cell cycle progression, we found that the CDK inhibitor gene, *KIP-RELATE PROTEIN1* (*KRP1/ICK1)* (*AT2G23430*), is the sole cell-cycle gene directly bound and downregulated by MUTE (Fig. 1C,D, Table S1).

### Dynamics of MUTE accumulation during the single terminal division event

Because our survey of MUTE direct targets identified cell cycle genes throughout the cell cycle (Fig. 1), we sought to address the dynamics of MUTE protein accumulation in the specific context of the single terminal division of a GMC. It has been shown that functional MUTE-GFP proteins become detectable shortly after the last asymmetric cell division of amplifying meristemoids (Han et al. 2018). To simultaneously monitor the dynamics of MUTE protein accumulation and cell-cycle phase, we generated transgenic Arabidopsis plants expressing both *MUTEpro::MUTE-GFP* and the multi-color cell cycle indicator PlaCCI (Desvoyes et al. 2020) (Fig 1A, Supplementary Figs. S1A and B). PlaCCI is comprised of three fluorescent-protein-tagged markers, CDT1a-eCPF, which marks G1 phase, HTR13-mCherry, which is notable in S-G2-M phase, and NCYCB1;1-YFP, which marks mitotic events (Desvoyes et al. 2020)(See Fig. 1B, Supplementary Fig. S1A)

Our long-term time-lapse live imaging of germinating cotyledon epidermis revealed two distinct types of GMCs. The first type do not exhibit any sign of asymmetric amplifying divisions and immediately proceed to the GMC division upon germination (Supplementary Fig. S2A, Supplementary Video S1). We classified them as pre-formed GMCs during embryogenesis, and omitted them from a further analysis. The second type exhibit a typical amplifying asymmetric division, indicative of their initial identity as meristemoid mother cells, and then execute a terminal, single SCD of GMCs to form a pair of guard cells (Fig. 2, Supplementary Fig. S2B, Supplementary Video S2). We therefore categorized these cells as those that underwent post-embryonic *de novo* progression of stomatal-lineage development (‘post-embryonic *de novo* GMCs’).

**Fig. 2:**
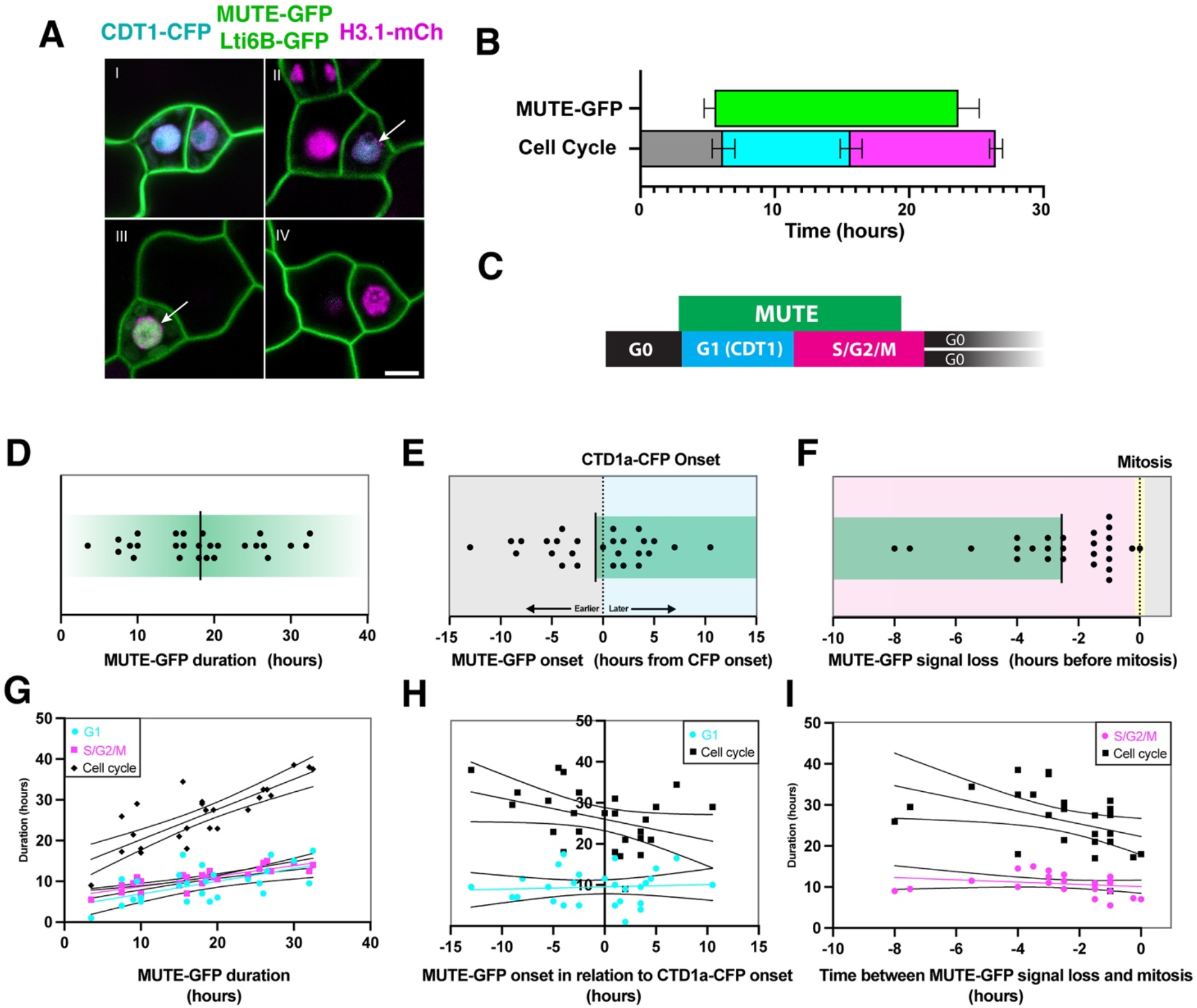
MUTE-GFP expression and cell cycle phase correlation during the terminal stomatal symmetric division. (A) Single still images of 3-day-old Col-0 cotyledon expressing MUTE-GFP, cell cycle marker PlaCCI and cell membrane marker Lti6B-GFP. Note that MUTE-GFP expression turns on during G1 phase (CDT1a-CFP, arrow point to nucleus) and vanishes during S/G2 phase (H3.1-mCherry). (B-I) Duration and timing of MUTE-GFP expression and PlaCCI cell cycle phases during stomatal symmetric divisions from time-course confocal images of 3-4dag Col-0 cotyledons. (B) Measured MUTE-GFP and PLaCCI cell cycle phases. (C) Schematic of expression windows from B. (D) MUTE-GFP duration. (E) Time between CDT1a-CFP appearance and MUTE-GFP appearance. (F) Time between MUTE-GFP signal loss and mitosis. (G) Correlation between MUTE-GFP window duration and cell cycle length. (H) Correlation between values in panel E and cell cycle length. (I) Correlation between values in panel F and cell cycle length. Solid lines represent linear regression, dashed (curved) lines represent 95% confidence level of linear regression. n=27 from 3 samples for all. P values: 0.05 > * > 0.005, 0.005 > ** > 0.0005, 0.0005 > *** > 0.00005, 0.0005 > **** Values and correlation statistics listed in Supplementary Dataset S1.

For these cells fully undergoing stomatal cell-state transitions, MUTE-GFP accumulation became evident shortly (4.8 ± 3.6 h) after the last ACD, 0.7 ± 5.4 hours prior to CDT1-eCFP (cyan) appearance (Fig. 2B,C,E). The MUTE-GFP signals persisted through G1, S and G2 phases, lasting an average of 18.2 ± 7.9 hours (Figs. 2B,C,D), then disappeared an average of 2.5 ± 2.0 hours before the M phase marked by NCYCB1;1-YFP (yellow) (Fig. 2B,C,F). The duration and onset time of MUTE-GFP correlate well with cell cycle length (Fig. 2G,H), suggesting that MUTE and cell cycle regulators are co-regulated. Taken together, our analysis revealed the presence of two classifications of GMCs, embryonic and post-embryonic, both undergoing the single terminal division. More importantly, the simultaneous time-lapse imaging of MUTE and PlaCCI revealed that MUTE accumulation initiates immediately after the last ACD of a meristemoid, sustains through G1-S-G2, and disappears prior to the M phase, consistent with the finding that MUTE transcriptionally induces cell cycle genes which function through each of the cell cycle phases (Fig. 1).

### FAMA turns on in the late G2 phase before the terminal division

It has been shown that MUTE directly induces *FAMA* expresion. This in turn creates a regulatory network motif that can generate a single pulse of cyclin/CDK expression during the terminal division (Han et al. 2018). To examine in which cell cycle phase FAMA accumulates, we generated transgenic Arabidopsis plants expressing both *FAMApro::FAMA-GFP* and the cell cycle indicator PlaCCI (Fig. 3A). Time-lapse imaging analyses show that FAMA-GFP accumulation begins in the late G2 phase, an average of 2.9 ± 1.6 hours prior to the terminal GMC division (Fig. 3A-C,D), closely overlapping with the end of the MUTE-GFP expression window (Fig. 3D). This is consistent with the previous inducible *MUTE* study, which showed delayed induction of FAMA at 8 hours after the induced *MUTE* overexpression (Han et al. 2018). The duration of FAMA-GFP accumulation is nearly identical to that of MUTE-GFP, lasting 18.3 ± 3.9 hours on average (Fig. 3D). FAMA-GFP persists into G0 of the sister guard cells, terminating 14.5 ± 3.7 hours after mitosis (Fig. 3F).

**Fig. 3:**
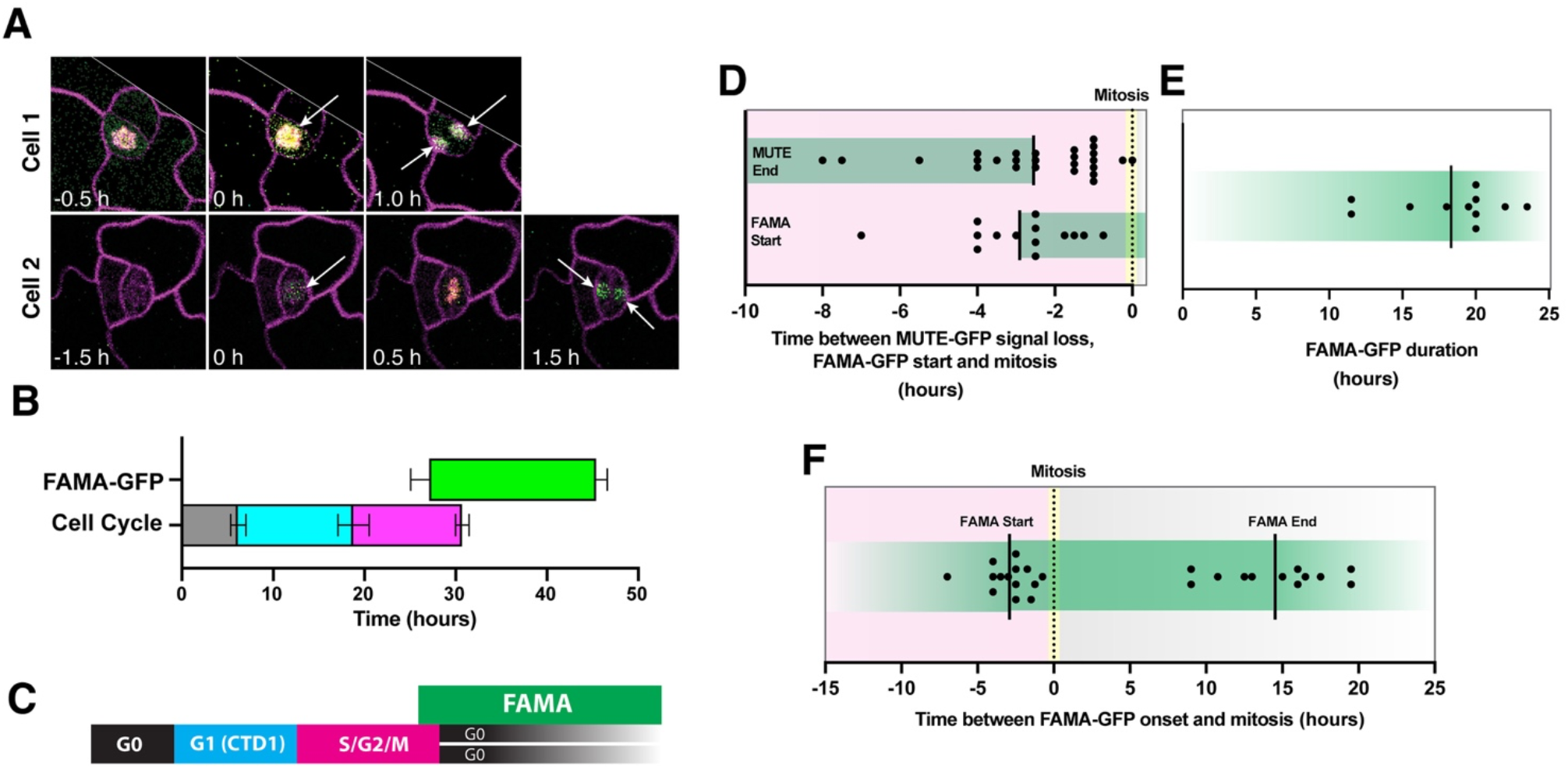
FAMA-GFP expression during the terminal stomatal symmetric division. (A) Representative timelapse confocal imaging of 3-day old cotyledon expressing FAMA-GFP, cell cycle marker PlaCI and cell membrane marker PM-RB. White arrows indicate FAMA-GFP expression before and after mitosis. (B-F) Duration and timing of FAMA-GFP expression and PlaCCI cell cycle phases during stomatal symmetric divisions from timecourse confocal images of 3-4dag Col-0 cotyledons. (B) Measured FAMA-GFP and PLaCCI cell cycle phases. (C) Schematic of expression windows from B. (D) Time between FAMA-GFP appearance and mitosis, compared to the time between MUTE-GFP disappearance and mitosis (from panel 2F). (E) Duration of FAMA-GFP expression (F) Time between FAMA-GFP appearance and mitosis (left); time between FAMA-GFP disappearance and mitosis (right). n = 14 from 3 samples

### Meristemoids that cannot commit to differentiation elongate G1 as reiterating ACDs

In the Arabidopsis developing cotyledon and leaf epidermis, meristemoids typically undergo ∼3 proliferative, ACD prior to terminally differentiating into a GMC under the control of MUTE (Han and Torii 2016; Nadeau and Sack 2002). In the absence of *MUTE*, the meristemoids continue to asymmetrically divide, reduce in size, and eventually arrest (Pillitteri et al. 2007). To understand the cell cycle behaviors of these *mute* meristemoids that are incapable for differentiation, we quantitatively analyzed the cell cycle dynamics of each round of ACDs in wild-type and *mute* meristemoids using time-lapse imaging (Fig. 4, Supplementary Figs. S1C, S2C, Supplementary Video S3. No significant differences in G0 or G1 duration were observed when comparing between sequential divisions in control plants, and only a slight difference was observed in S/G2/M duration between meristemoids undergoing division one compared to division three (Fig. 4A, B). On average, G0 (mitosis to CDT1a-CFP onset) lasted 8.5 ± 4.3 hours, G1 lasted 6.6 ± 5.0 hours, and the remaining three phases together (S, G2, and M) lasted 7.2 ± 2.2 hours, for a total cell cycle of 13.8 hours starting at G1 (using CDT1a-CFT loading as a proxy), consistent with the previous finding (Han et al. 2022), and the entire division cycle for 22.3 hours (Fig. 4B).

**Fig. 4:**
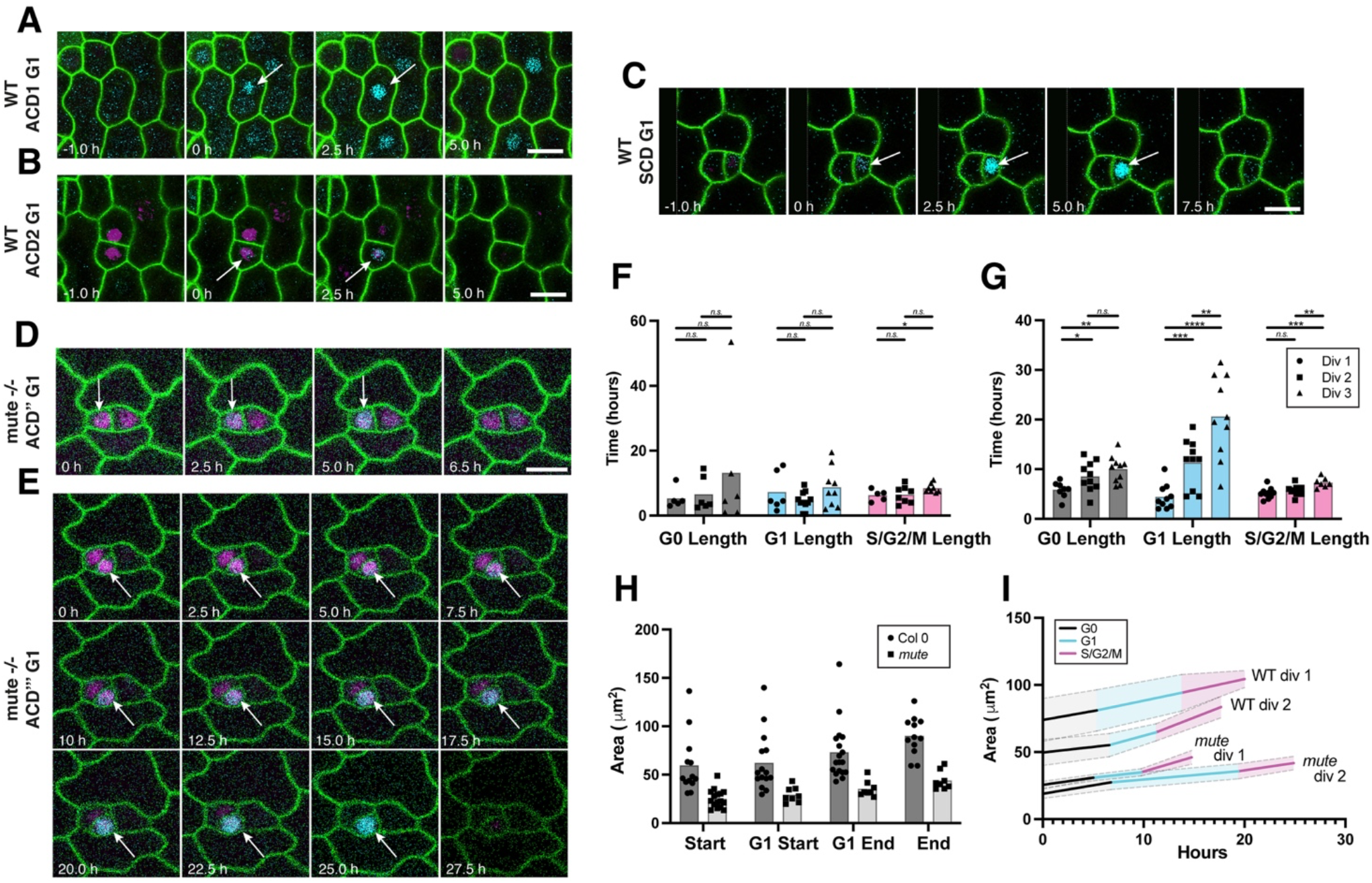
G1 phase is extended in reiterative ACDs in the absence of MUTE. (A-E) Representative timelapse confocal images of G1 phase of ACD1 (A), ACD2 (B) and SCD (C) in wild-type (WT), and ACD2(‘‘)(D), ACD3(‘‘‘)(E) in *mute* mutants from 1-to 5-day-old cotyledons expressing both PlaCCI and Lti6B (green). White arrows indicate the nuclei with CDT1-CFP fluorescence, representing G1. Scale bar, 10um. (F, G) Comparison of individual cell cycle length of three rounds of ACD between WT (F) and *mute* mutants (G). For WT, div1 n = 7; div2 n = 11, div3 n = 13, samples = 6. For *mute*, div1 n = 15; div2 n = 15, div3 n = 15, samples = 2 (H) Area of control and *mute* meristemoids at various stages of the cell cycle. For each cell, area was measured just after a new cell wall formed (Start), at CTD1a-CFP onset (G1 Start), at CTD1a-CFP disappearance (G1 end) and just prior to cytokinesis (End). Control n = 20, *mute* n = 18. Wild-type (WT) samples = 6, *mute samples =2* (I) Growth through each phase of the cell cycle using individual area values from panel H and average values for phase length from panels F and G. Control div 1 n = 8, div 2 n = 5, *mute* div 1 n =4, div2 n = 4. Wild-type (WT) samples = 6, *mute* samples *=* 2 Scale bars = 10 µm. For pairwise comparisons, two-tailed students t-test was used. P values: 0.05 > * > 0.005, 0.005 > ** > 0.0005, 0.0005 > *** > 0.00005, 0.0005 > ****

Interestingly, however, we observed an elongation of the ACD cell cycle over time in *mute* meristemoids but not in the wild-type meristemoids. Lengthening of each phase of the cell cycle with successive meristemoid divisions is accompanied with significant extension of the G1 phase. As shown in Fig. 4C, D, the avarage G1-phase duration was increased by roughly 5-fold, from 4.2 ± 2.4 hours in the first division to 22.4 ± 7.6 hours in by the third division. Next, to ascertain if G1, specifically, is being lengthened to allow small *mute* meristemoids to expand to a required division threshold, we measured the cellular area of the meristemoids from wild-type and *mute* mutants throughout the cell cycle in each round of ACDs. We found that cellular growth of meristmoids is not restricted to G1, but is in fact relatively uniform (Fig. 4E,F,G). Rather than driving the cellular growth of meristemoids for futher ACDs, we conclude that the gradual G1-phase extension during ACDs of *mute* meristemoids likely reflects a stepwise progression toward developmental arrest, a phenotype that has been observed in aged, uncommitted meristemoids (Pillitteri et al. 2008).

## DISCUSSION

In this study, we investigated the role of MUTE in governing cell cycle during the terminal division cycle of a stomata, and whether MUTE expression itself is reciprocally gated by the cell cycle. Furthermore, we revealed that the loss of *MUTE* has phase-specific impacts on cell cycle dynamics in meristemoids, which are not able to switch from a proliferative mode to differentiation. From comparative MUTE Chip-Seq and induced MUTE RNA-Seq analyses, we have shown that direct MUTE targets extend from G1 (e.g. *CYCD5;1*) (Han et al. 2018) to the G1/S checkpoint, S-phase, and G2 to G2/M checkpoint, indicating that MUTE is capable of installing a module of cell-cycle machinery that is sufficient to execute a complete division cycle. The installation of a complete MUTE-derived cell cycle module has important implications – it may provide opportunities for MUTE-SCD-specific regulatory input by incorporating specific cell cycle componenets into the developmental network logic. Importantly, the direct upregulation of checkpoint control components by MUTE, including G1/S transcription factors E2FF/DEL3 and E2F2, S-phase DNA helicase MCM3, and M-phase APC/C regulator GIG1 likely ensures that SCD is executed once MUTE is expressed above certain threshold. In this regard, it is worth mentioning that ectopic MUTE overexpression could induce SCD in epidermal cells with a pavement cell-like character, cells that would otherwise be division incompetent (Han et al. 2018; Pillitteri et al. 2008; Pillitteri et al. 2007).

Using the cell cycle marker PlaCCI in conjunction with MUTE-GFP, we found that MUTE-GFP signals always overlap with the G1 and S/G2 phases and are confined to a single division cycle (Fig. 2). This restricted accumulation window may reflect feedback from cell-cycle machinery, or simply be a function of protein stability and non-cell-cycle network logic. However, our data also show that MUTE duration correlates with cell cycle length (Fig. 2G), and that MUTE-GFP accumulation always diminishes prior to mitosis (Fig. 2F, Supplementary Fig. S2B,), suggesting the possibility that MUTE protein is regulated, at least in part, by cell-cycle machinery. To date, no studies have identified promoter elements or protein motifs in *MUTE* that resemble known cell cycle regulatory mechanisms, although a peak of H3K27me3 at the MUTE locus (Lee et al. 2019) could be an opportunity for cell-cycle-dependent RBR1-based chromatin remodeling. Future studies to manipulate the cell cycle lengths and addressing the effects on MUTE accumulation duration, may inform whether MUTE expression itself is under the cell cycle control.

We found that FAMA, a known target of MUTE, begins to accumulate several hours prior to the symmetric cell division at the late G2 phase, coinciding with the timing of MUTE protein disappearance (Fig. 3D). The rapid disappearance of MUTE concurrent to FAMA appearance implies a possible mutually-exclusive nature of regulation. Indeed, MUTE-induced cell cycle machinery, including *CYCD5;1* and and *CDKB1;1*, persists in the GMC in the absence of *FAMA*, triggering multinumeral divisions in *fama* GMCs (Han et al. 2018; Ohashi-Ito and Bergmann 2006). This *fama* phenotype is consistent with the notion that MUTE remains active in the absence of *FAMA*. While the actual, molecular mechanism of this mutual-exclusiveness is unknown, it likely involves an epigenetic mechanism. Indeed, as the seedling gets older, the *MUTE* locus becomes epigenetically repressed with the deposition of repressive chromatin marks, H3K27me3, which may be limiting the developmental windows of *MUTE*-expression (Lee et al. 2014). FAMA can act as an epigenetic regulator of stomatal lineage genes, including *SPCH* (Lee et al. 2019). Alternatively, sequential expression of MUTE and FAMA may reflect their expression windows within the cell cycle, mimicking a MUTE-FAMA mutual-exclusion relationship through opposing cell-cycle inputs during G2. It would be an interesting future direction to investigate the intersection of G2-specific transcriptional regulation (e.g. MYBR3s) (Kobayashi et al. 2015) on the expression of MUTE and FAMA.

Previous studies have shown that the cell cycle within the stomatal-cell lineage is pliable (Han et al. 2022). Depending on the presence or concentration of CKIs such as the MUTE-induced SMR4, delecerated cell cycle can impact cellular differentiation and fate segregation. We found that, as meristemoids continue to divide in the absence of *MUTE*, their G1 phase becomes extended in successive divisions. This is different from the MUTE-orchestrated deleratation of cell cycle in SCD, given that different sets of core cell cycle regulators are operating (Han et al. 2022). Cell cycle arrest in *mute* meristemoids could be a consequence of cell size being a limiting factor to support mitotic division. Based on such a hypothesis, the observed specific G1-phase extention implies that G1 period might be utilized for cellular growth. However, our observations that growth occurs in a relatively uniform manner throughout the cell cycle (Fig. 4H,I) contradicts this hypothesis. Our additional observations, as well as those of other studies in mammals (He et al. 2009; Zetterberg and Larsson 1985), suggest that G1 may simply be the most pliable/dynamic phase, and G1 extension may represent a strategy to increase the cell cycle duration overall.

We propose that, in meristemoids, G1 phase operates as a flexible GO-NO-GO threshold, after which the other phases of the cell cycle proceed at a nearly constant rate, until mitosis. Since MUTE protein accumulation starts in G1 (Fig. 2), it is also conceivable that G1 lengthening in the absence of MUTE is a survival strategy – providing a larger activation window for MUTE in aging meristemoids. In contrast to the fate-mixing and trans-fating observed due to the early stomatal-cell lineage expression of SMR4 induced G1 elongation (Han et al. 2022), *mute* meristemoids retain proflierative status presumably due to extended action of the MUTE’s predecessor *SPCH*, which promotes mitotic potential of the meristemoids (Lau et al. 2014). This suggests that cell cycle alteration alone is not sufficient to cause fate mixing of stomatal guard cells and non-stomatal pavement cells in the absence of *MUTE*. Future studies of how cell cycle machinery shapes the expression of MUTE and other stomatal master regulatory transcription factors will illuminate our understanding of specialized cell-type differentiation in plants.

## Materials and Methods

### Plant Materials

*Arabidopsis* accession Columbia (Col-0) was used as wild-type. The following mutants/transgenic lines have been published elsewhere: *MUTEpro::MUTE-GFP* (Pillitteri et al. 2007); *MUTEpro:::MUTE-tagRFP* (Qi et al. 2017); *mute-2*, used as *mute* mutant (Pillitteri et al. 2008); *FAMApro::FAMA-GFP* (Han et al. 2018); Lti6b (Kurup et al. 2005); PlaCCI (Desvoyes et al. 2020). Higher-order mutant/marker lines were generated by genetic crosses and their genotypes were confirmed. Sterilized Arabidopsis seeds were grown on half strength of Murashige and Skoog (MS) media with 1% sucrose and stratified for 2-3 days at 4°C. The seedlings were grown at 22°C under long-day condition, and 10∼14-day-old seedlings grown on MS media were transplanted to soil to harvest seeds. For the selection of *mute-2/+*, seeds were grown on the 1/2 MS media supplemented with 50 µm/L Kanamycin (Fisher, BP906-5).

### Confocal microscopy and time-lapse imaging

Confocal microscopy images were either acquired using Stellaris-8 FALCON (Leica) using a 63x oil-lens for high resolution imaging and 40x oil-lens for data acquisition or SP5-WLL/Argon (Leica) using a 63x water-lens for high resolution imaging and 20x dry-lens for data acquisition. The time-lapse imaging of germinating cotyledons expressing PlaCCI (Desvoyes et al., 2020) in WT and *mute*, or together with MUTE-GFP and FAMA-GFP was performed as described previously (Han et al. 2022; Peterson and Torii 2012). 1-day old germinated seedlings were dissected from seeds and placed onto chamber slides (Thermo Fisher Scientific, Nunc Lab-Tek II #155379), which were then placed on a motorized stage. Leica Stellaris 8 FALCON with the following conditions: CFP, excitation at 458 nm and emission from 464 to 510 nm; YFP excitation at 514 nm and emission from 520 to 560 nm; mCherry and tagRFP, excitation at 561 nm and emission from 570 to 620 nm. Signals were visualized sequentially using separate HyD detectors (HyDX/HyDS) in TauSeparation mode. Leica SP5 images were imaged using SP5-WLL/Argon with the following conditions: CFP, excitation at 458 nm and emission from 468 to 600 nm; GFP, excitation at 488 nm and emission from 490 to 546 nm; YFP, excitation at 514 nm and emission from 524 to 555 nm; mCherry, excitation at 560 nm and emission from 565 to 620 nm. The time-lapses were collected at 30-min intervals using a 63x oil-lens, zoom factor 1.3-1.5 for high resolution images and at 30-min intervals using a 40x oil-lens or 20x dry-lens, zoom factor 1 for data acquisition

### Bioinformatic analysis and data visualization

For extracting the cell cycle genes from our MUTE ChIP-seq and RNA-seq data (Han et al. 2022; Han et al. 2018), cell cycle related genes were extracted from GO:0007049, GO:0051726, GO:0051321, GO:0007346, GO:0007050, GO:0000082, GO:0010389, GO:0010971, GO:0000086, GO:0045787, GO:0071158, GO:0000278, GO:0006267, GO:0007093, GO:0045786, GO:0045931, GO:0051446, GO:0007113, GO:0044843, GO:0045930, GO:0051445, GO:0060154, GO:0060184, GO:1900087, GO:1902749, GO:0090266 and were combined with manually curated genes. Genes increased and/or decreased by MUTE more than Log 2FC (Fold change) 0.4 and targeted by MUTE were extracted. The data obtained were visualized using R-package “Venn Diagram”. Bedgraph file from previous MUTE ChIP-seq was generated and visualized in IGV browser (ver. 2.4.11) (Robinson et al. 2011).

### Image processing and quantitative analysis

A series of either Z-stack confocal images (time-lapse imaging) or single plane images were obtained to capture fluorescent protein signals (CFP, GFP, YFP, and mCherry). Raw data were collected with 512 × 512 pixel image and imported into Fiji-ImageJ v1.8.0_66 to generate RGB images/z-stacks using the channel merge function. To correct for drift of multichannel z-stacks the “StackReg” plugin was applied. Movies are played with 7fps. The area of meristemoids were quantified with FIJI-ImageJ v2.3. Statistical analyses were performed using Graphpad Prism v9.2. For two-sample comparisons, student t-tests were performed. Graphs were generated using Graphpad Prism v9.2. The value of n, the number of each experiment or samples, and how statistical significance was defined are indicated in each relevant figure legend.

## ACKNOWLEDGEMENTS

We thank Dr. Soon-Ki Han, Prof. Crisanto Guiterrez and Dr. Bénédicte Desvoyes for the original MUTE ChIP-seq data and PlaCCI construct, respectively, and Hyemin Seo for assisting genotyping.

## AUTHOR CONTRIBUTIONS

Conceptualization, K.U.T.; Experimental Design, D.T.Z, A.H., K.U.T.; Performance of experiments, D.T.Z., A.H., K.U.T.; Bioinformatics Analysis, E.-D.K., K.U.T.; Visualization, D.T.Z., A.H., K.U.T.; Writing-Original Draft, K.U.T., D.T.Z.; Writing-Review & Editing, all authors; Project Administration, K.U.T.; Funding Acquisition, K.U.T.

